# Dynamics of Macrophage Polarization Reveal Unique Phenotypes During Mouse Postpartum Uterine Remodeling

**DOI:** 10.1101/2025.09.19.676950

**Authors:** Maria Natalia Chaves Rivera, Sydney Bell, Abigail Solomon, Harvey J. Kliman, Aya Tal, Reshef Tal

## Abstract

**Background:** Macrophages play essential roles in uterine physiology during pregnancy and the postpartum (PP) period. In various organ systems, macrophages exhibit distinct M1 (pro-inflammatory) and M2 (anti-inflammatory) polarization states, the balance of which is key not only in scarless wound healing but also in pathological fibrosis. While macrophages are central players in PP uterine remodeling, their polarization dynamics during this period and associated spatial distribution remain largely unknown.

**Methods:** Uterine tissues were collected from C57BL/6J wild-type mice at various PP days (PPD 1, 3, 7, 14, 21, 28) and non-pregnant controls. Multicolor flow cytometry was used to analyze immune populations and macrophage polarization states. Macrophage polarization was determined using CD86 (M1 marker) and CD206 (M2 marker). Immunofluorescence (IF) and immunohistochemistry (IHC) were used to examine the colocalization of M1 and/or M2 markers alongside F4/80 to assess macrophage polarization status and tissue distribution.

**Results:** On PPD1, uterine macrophage numbers were higher compared to all other time points, with significantly increased M2 macrophages (CD86-CD206+) (43% p<0.0001) compared to PPD3 (5.1%), PPD7 (2.4%), PPD14 (2.3%), and non-pregnant (1.9%). A rapid polarization shift to M1 macrophages (CD86+CD206-) occurred on PPD3 (50.6%), PPD7 (66.5%), and PPD14 (58.8%) compared to PPD1 (11.9% p<0.0001). Additionally, there was a trend of increased M2 macrophages on PPD21 (10%) and PPD28 (5%). We observed a unique macrophage subset of mixed phenotype (co-expressing CD86+CD206+) significantly higher on PPD 1 (37.2% p<0.05) compared to NP (7.8%), PPD7 (11.6%), and PPD14 (26.4%). IF co-staining confirmed these dynamics, showing the highest percentage of macrophages co-expressing CD206+F4/80+ on PPD1 (77.2%), significantly higher than all other time points. IHC revealed the specific temporal and spatial distribution of F4/80+ and CD206+ cells within the uterine endometrial stroma adjacent to the myometrial junction in the early PP period.

**Conclusions:** The predominance of M2-polarized macrophages on PPD1 followed by a rapid shift to M1 phenotype from PPD3 to PPD14 indicates an evolving inflammatory response during the early postpartum period. Repolarization to M2 profile on PPD21, along with increased abundance of a mixed population (M1/M2), uncovers the dynamic complexity of macrophage polarization during PP uterine remodeling, providing new insights into scarless uterine tissue repair.

## INTRODUCTION

The female reproductive system undergoes profound structural and functional adaptations during pregnancy to support fetal development and childbirth. Notably, the uterus expands up to 20 times its normal size in humans, primarily due to an increase in both the number and size of the myometrial cells. Following delivery, it gradually returns to its baseline non-pregnant state within 42 to 56 days.^1,2^ This postpartum remodeling process, also known as uterine involution, involves extensive cellular and molecular changes that restore the uterus to its pre-pregnancy dimensions and function. It includes the regeneration of the endometrial lining, reduction in uterine vascularity, and decrease in myometrial mass. These changes are primarily driven by alterations in collagen, elastin, and smooth muscle composition, coupled with shifts in hormonal and inflammatory signaling.^3,4^ This remodeling is largely facilitated by elevated activity of enzymes such as matrix metalloproteinases (MMPs), elastase, and cathepsin B.^5–7^ These enzymes are produced and regulated by various cell types, including fibroblasts, smooth muscle cells, and macrophages, which collectively orchestrate the structural remodeling of the uterus.^6–8^

The postpartum uterus is characterized by a dynamic immune environment, with strong activation of innate immunity and regulated adaptive responses that are crucial for bacterial clearance, tissue repair, and preparation for future fertility. After delivery, the uterus transitions from an immune-tolerant state required for pregnancy to a heightened inflammatory environment that supports repair. Neutrophils and macrophages are swiftly mobilized to the uterus, serving as key early responders. Together with dendritic cells and endometrial epithelial cells, which provide barrier protection, they coordinate both the inflammatory response and subsequent tissue repair and remodeling.^9–11^

Macrophages play a central role in postpartum uterine remodeling and tissue repair. Following delivery, these immune cells are actively recruited to the uterus, where they exhibit heightened activation and phenotypic plasticity to meet the dynamic demands of the healing environment.^12^ One of their key functions is the degradation of excess extracellular matrix components, including collagen, as well as the clearance of cellular debris, both of which are essential for supporting uterine involution.^13^ In addition to these roles, macrophages facilitate the elimination of senescent cells from the implantation fibrotic scar area, a process crucial for restoring normal uterine architecture and function.^14^

Macrophages exhibit remarkable functional plasticity, adopting distinct polarization states shaped by their developmental origin and the surrounding tissue microenvironment. Traditionally, they are classified under the M1/M2 paradigm: classically activated (M1) macrophages, which are pro-inflammatory, and alternatively activated (M2) macrophages, which are anti-inflammatory and involved in tissue repair.^15,16^ In the uterus, this dynamic polarization is tightly regulated and plays critical roles throughout the reproductive cycle. During early pregnancy, macrophages support implantation and vascular remodeling by displaying a mixture of M1-and M2-associated features, while later stages are dominated by M2 macrophages that promote fetal tolerance. At parturition, a shift back toward proinflammatory M1 macrophages facilitates uterine contractions and placental separation.^12^ Although M2 macrophages are thought to contribute to immediate postpartum repair, their distribution, polarization state dynamics, and functions across the entire postpartum period remain poorly defined.^12,14^

The conventional M1/M2 classification, however, is now recognized as overly simplistic and insufficient to capture the full spectrum of macrophage activation states.

Studies in diverse tissues have shown that macrophages can transition between phenotypes in response to environmental cues. During tissue repair, for instance, macrophages often shift from a pro-inflammatory (M1-like) to an anti-inflammatory (M2-like) profile.^17^ Moreover, macrophages can co-express markers traditionally associated with both M1 and M2 phenotypes,^18,19^ and transcriptomic studies indicate that polarized macrophages can be reprogrammed to the opposite phenotype without retaining a “memory” of their prior activation.^20^

Given this complexity and the lack of detailed profiling of macrophages in the postpartum uterus, we sought to characterize the polarization states and temporal dynamics of uterine macrophages throughout the postpartum period. Increased understanding of macrophage dynamics in this context may provide key insights into the mechanisms that enable scarless postpartum uterine healing.

## MATERIALS AND METHODS

### Animals

C57BL/6J wild-type mice were obtained from Jackson Laboratories (Bar Harbor, Maine) and maintained under an approved Yale University Institutional Animal Care and Use Committee (IACUC) protocol (2023-20496) at the Yale Animal Resource Center. Mice were housed in a controlled environment with a 12-hour light/dark cycle and had ad libitum access to food and water. Ten-twelve weeks old female mice were bred with wild-type fertile males at a 1:3 male-to-female ratio. Females were weighed every other day to detect pregnancy-related weight increase. Females were separated from males upon >8g weight increase from baseline and monitored daily until delivery. Postpartum mice were euthanized at the following postpartum days (PPD) for analysis: PPD1, PPD3, PPD7, PPD14, PPD21, and PPD28. Additionally, cycling non-pregnant (NP) mice were euthanized in the diestrus phase as controls. The bottom third of the left uterine horn of each mouse was fixed in 4% paraformaldehyde (PFA) for histopathological analysis, while the remaining tissue was used for multicolor flow cytometry (Figure 1).

**Figure 1.**
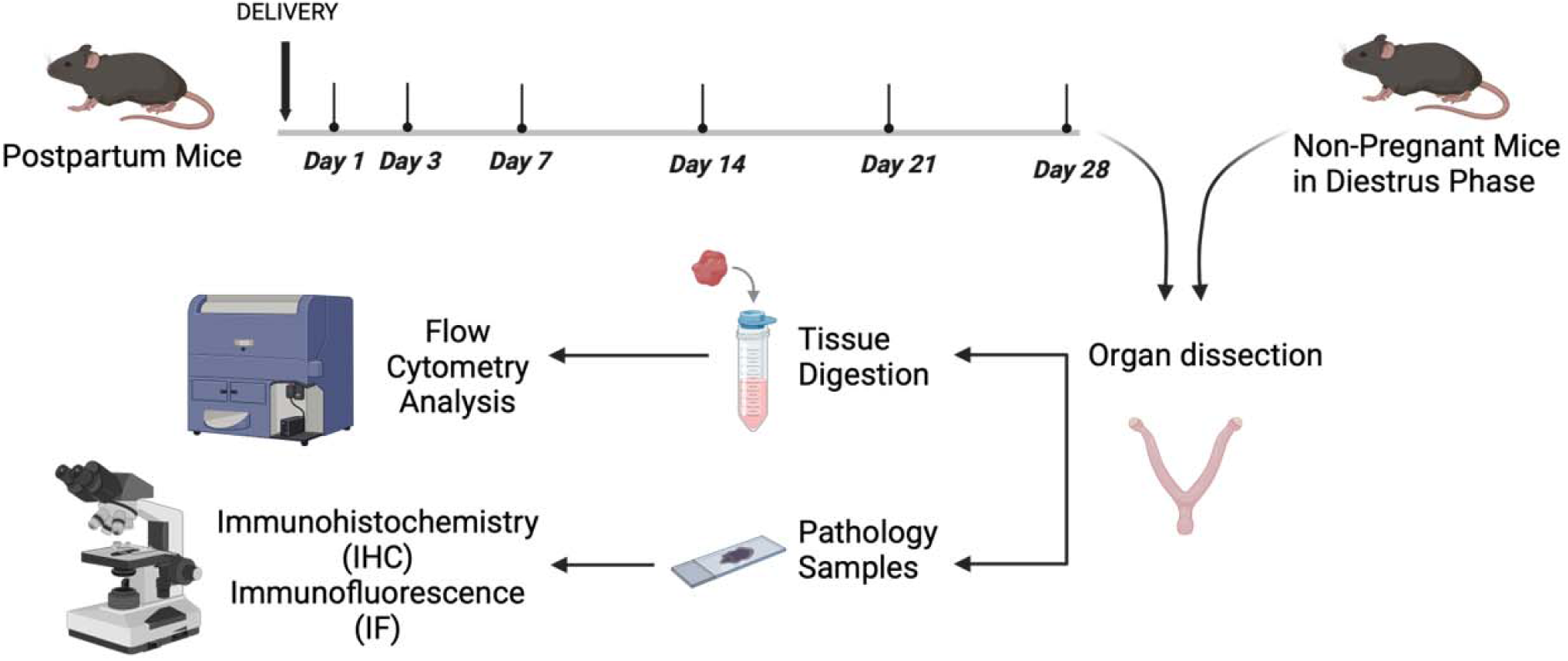
Schematic of the experimental design. Wild-type (WT) C57BL/6J female mice were sacrificed at various postpartum days (PPD 1, 3, 7, 14, 21, and 28) (n = 5-6/group). Cycling non-pregnant (NP) controls (n = 5) were sacrificed during the diestrus phase. Uteri were dissected and processed for flow cytometry, as well as for histopathological analysis, including immunofluorescence and immunohistochemistry.

### Vaginal cytology

Estrous cycling in virgin non-pregnant mice was induced by transferring the females to a cage containing bedding from previously-housed males. Vaginal cytology was performed daily to monitor estrous cycle stages as previously described.^21^ Briefly, a sterile pipette was used to inject and withdraw 20 μL of PBS within the vaginal canal to collect vaginal cells. The collected cells were smeared onto a slide, mounted with a coverslip, and examined under a light microscope.

### Flow Cytometric Analysis

Following uterine extraction from non-pregnant or postpartum mice at various time points, tissues were immediately placed in Hank’s Balanced Salt Solution (HBSS) medium on ice. The tissue was finely minced, followed by enzymatic digestion in HBSS containing 1 mg/mL Collagenase B (Roche Diagnostics) and 0.1 mg/mL deoxyribonuclease I (Sigma-Aldrich) for 30 minutes at 37°C, with thorough mixing every 10 minutes. The resulting cell suspension was filtered through a 70 μm mesh, centrifuged at 2000 rpm for 8 minutes at 4°C, washed in FACS buffer (2% FBS in phosphate-buffered saline (PBS)), and centrifuged again at 1500 rpm for 5 minutes at 4°C. The cell pellet was resuspended in 100 uL FACS buffer for antibody staining and blocked for 10 minutes using 2 μL TruStain fcX anti-mouse CD16/32 (#101320, BioLegend). Staining with a specific antibody cocktail (Supplemental Table S1) was done on ice for 30 minutes. After staining, cells were washed twice with FACS buffer, resuspended in PBS, and analyzed on a BD LSR Fortessa Cell Analyzer Flow Cytometer (Yale Flow Cytometry Core, New Haven, CT). Live gates were applied to forward and side scatter plots to exclude debris and nonviable cells. Unstained and IgG isotype controls were used for compensation and gating. Data was analyzed using FlowJo V10 software.

### Immunohistochemistry (IHC) and immunofluorescence (IF)

Uterine tissues were fixed in 4% PFA for 24 hours and then transferred to 70% ethanol. The tissues were embedded in paraffin, cut into 5 μm sections and mounted on slides at the Yale Pathology Tissue Services. Sections were deparaffinized with xylene and rehydrated through a graded ethanol series before undergoing heat-mediated antigen retrieval in 10 mM sodium citrate buffer (pH 6.0). Subsequently, slides were washed in PBS and incubated with hydrogen peroxide (H2O2) to block endogenous peroxidase activity. Blocking was performed for 1 hour at room temperature using 5% serum in PBS-T. Slides were incubated overnight at 4°C with rabbit anti-F4/80 primary antibody (1:500, #ab300421, Abcam, Cambridge, MA) or goat anti-CD206 primary antibody (1:400, #AF2535, R&D Systems, Minneapolis, MN). The following day, IHC sections were washed with PBS and incubated for 1 hour at room temperature with the appropriate biotinylated secondary antibody from ABC Vectastain Elite kit (1:200, #30014 or #30056, Vector Laboratories, Newark, CA). Detection was performed using ABC Vectastain Elite reagents with DAB and H2O2 (Vector Laboratories). Tissue sections were counterstained with hematoxylin staining solution (Ricca)and imaged using an ECHO Revolve microscope (Discover Echo, San Diego, California).

For immunofluorescence, rehydrated slides were incubated for 10 minutes in 0.1M ammonium chloride solution (pH 8.0) to block autofluorescence and permeabilized with PBST for 10 min. Antigen retrieval was performed as above. Sections were then blocked with 10% donkey serum in PBS or 1 hour, followed by overnight incubation with the same antibodies as above. The following day, slides were washed and incubated for 1 hour at room temperature with Alexa Fluor 594 donkey anti-rabbit (1:200, A21207, ThermoFisher Scientific, MA) and Alexa Fluor 488 donkey anti-goat (1:200, A11055, ThermoFisher Scientific, MA) secondary antibodies. Following incubation, slides were mounted using Vectashield fluorescent mounting media containing 4′,6-diamidino-2-phenylindole (DAPI) (Vector Laboratories) for nuclear staining. Negative controls included sections incubated without primary antibodies but with secondary antibodies. Images were acquired using an ECHO Revolve microscope (Discover Echo, San Diego, California) and analyzed with ECHO software.

### Statistical analysis

Data including *n*, mean, standard error, and statistical significance values, are indicated in the text or the figure legends. Statistical analysis was performed using Student’s *t* test (2-tailed), Mann-Whitney *U* test, or a 1-way ANOVA with multiple comparisons using the false discovery rate method as appropriate, using Prism 9.0 (GraphPad Software). *P* < 0.05 was considered to be statistically significant.

## RESULTS

### Macrophages exhibit distinct M1 and M2 polarization profiles throughout the postpartum period

To examine the polarization profiles and temporal distribution of M1 and M2 macrophages in the mouse uterus during the postpartum (PP) period, a time-course experiment was conducted in C57BL/6J wild-type mice. A schematic depicting the experimental workflow is provided in Figure 1. Mice were euthanized on postpartum days (PPD) 1, 3, 7, 14, 21, 28, or as non-pregnant (NP) virgin controls (n = 5–6/group). Multicolor flow cytometry was performed on uterine tissues collected from postpartum mice at designated time points and from cycling non-pregnant females in the diestrus phase, as confirmed by vaginal cytology. Macrophage polarization was assessed using M1 (CD86) and M2 (CD206) markers. These surface markers are commonly used to distinguish M1 and M2 macrophages, respectively.^22–24^ In each sample, the proportions of M1, M2, and M1/M2 co-expressing macrophages were quantified. The full gating strategy is illustrated in Figure 2A. Multicolor flow cytometry analysis revealed a marked increase in uterine leukocytes (CD45+) on PPD1, comprising 46.7% of total cells as compared to most of the later PP time points and NP mice (<25%, p<0.05) (Figure 2B). This early leukocyte influx was accompanied by a marked increase in granulocytes (Ly6G+) on PPD1 (Figure 2C), consistent with an acute inflammatory response during the immediate PP period. In contrast, the proportion of T cells (CD3+) remained unchanged across most of the time points, with a notable decrease on PPD28 (Figure 2D). Moreover, the proportion of uterine macrophages (CD11b+F4/80+) was highest on PPD1 compared to most other time points (Figure 2E). Within the uterine macrophage population (CD11b+F4/80+), a significant increase in M2-polarized macrophages (CD86-CD206+) was observed on PPD1, comprising approximately 43% of the total macrophages, followed by a rapid decline in this population reaching a nadir on PPD7 (Figure 3, A and C). The proportion of M2 on PPD1 was markedly higher than the one observed on PPD3 (5.1%, *p*<0.0001), PPD7 (2.4%, *p*<0.0001), PPD14 (2.3%, *p*<0.0001), and in NP controls (1.9%, *p*<0.0001).

**Figure 2.**
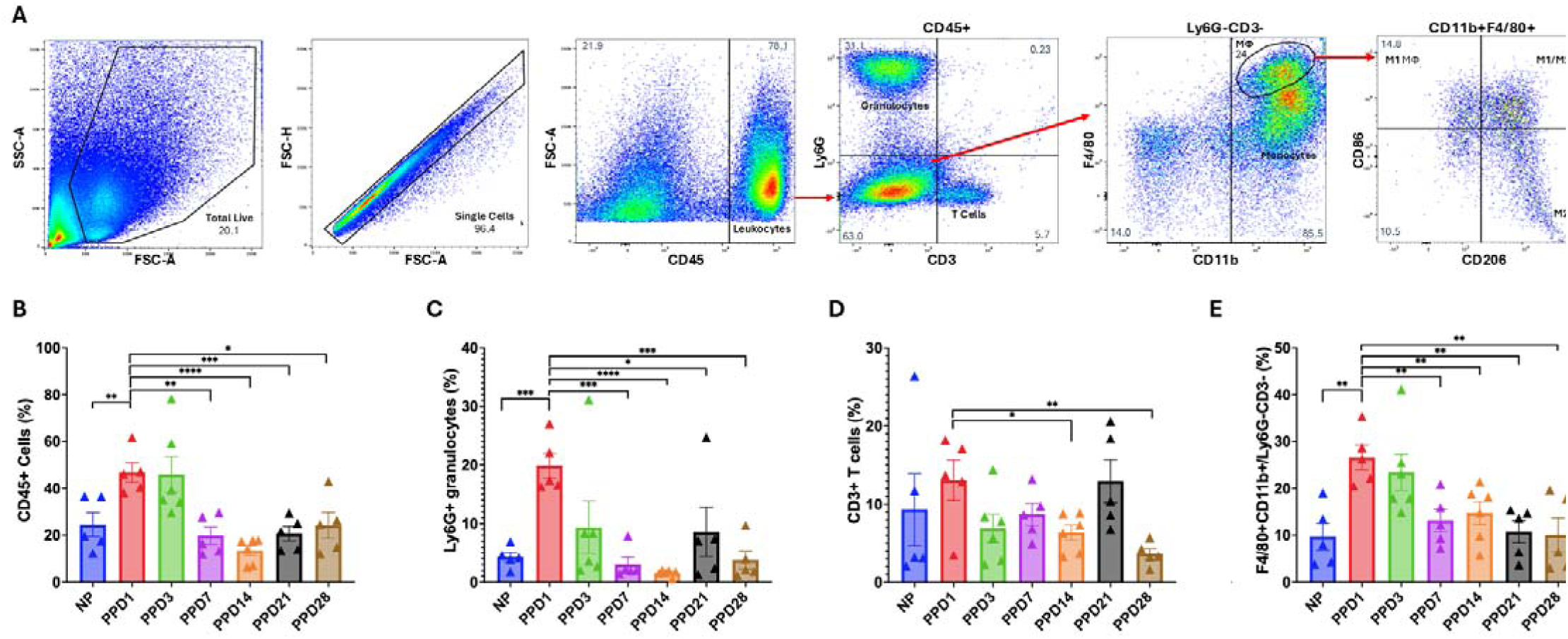
Gating strategy used for profiling uterine immune and specific macrophage populations in the postpartum period. (A) Multicolor flow cytometry was used to characterize immune cell populations and evaluate macrophage polarization in uterine tissues collected from non-pregnant (NP) diestrus mice and postpartum (PP) mice on days 1, 3, 7, 14, 21, and 28. The total live cell population was first selected by gating on viable cells and then excluding doublets, gating only single cells. Leukocytes were identified as CD45+ cells. Within this population, T cells and granulocytes were distinguished based on CD3 and Ly6G expression, respectively. CD3-Ly6G-cells were further analyzed, and monocytes/macrophages were gated as CD11b+. Macrophages were specifically identified as F4/80 high-positive cells within the CD11b+CD3-Ly6G-population. To evaluate macrophage polarization states, M1-like macrophages were assessed based on CD86 expression, while M2-like macrophages were identified using CD206 expression. (B) Percentage of CD45+ leukocytes out of gated live single cells. (C) Percentage of Ly6G+ granulocytes out of CD45+ leukocytes. (D) Percentage of CD3+ T cells out of CD45+ leukocytes. (E) Percentage of F4/80^high^CD11b+ macrophages out of CD45+CD3-Ly6G-cells. Data is presented as mean percentage ± SEM. *N* = 5-6 animals per group. *p < 0.05; **p < 0.01; ***p < 0.001; ****p < 0.0001.

**Figure 3.**
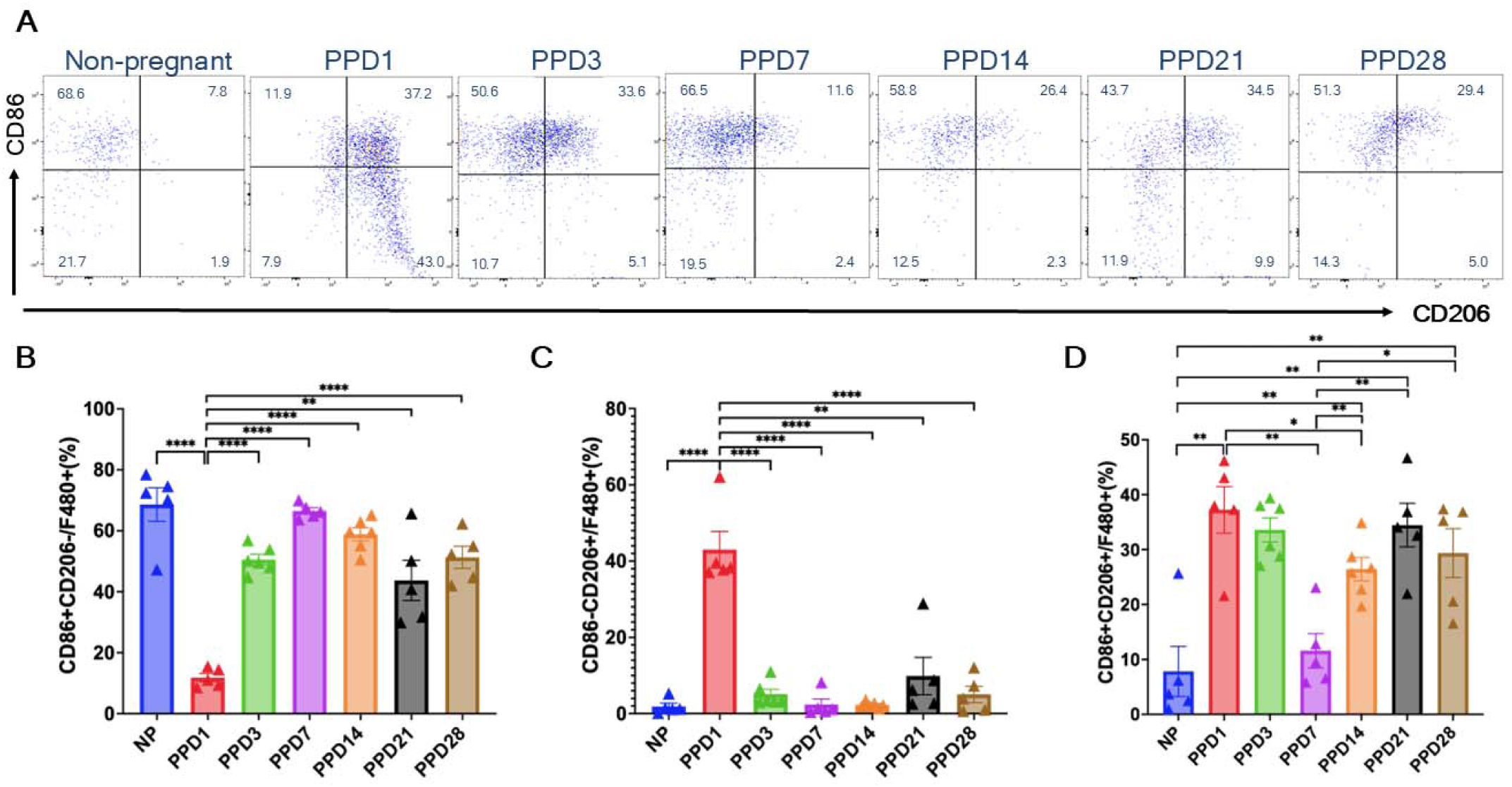
Uterine macrophage polarization during the mouse postpartum period. (A) Representative flow cytometry plots of F4/80^high^CD11b+ macrophages from uterine tissues in non-pregnant (NP) controls and postpartum (PP) mice at various time points. (B) Graphs showing proportions of M1 macrophages (CD86+CD206-), (C) M2 macrophages (CD86-CD206+), and (D) mixed macrophages co-expressing M1 and M2 markers (CD86+CD206+) across PP time points and in NP controls. Data is presented as mean ± SEM; *N* = 5-6 animals per group. \**p* < 0.05; **p < 0.01; \*\*\**p* < 0.001; ****p < 0.0001.

Concomitantly, a rapid shift toward the M1 phenotype (CD86+CD206-) was evident after PPD3, with M1-polarized macrophages (CD86+CD206-) comprising 50.6% of the macrophage population on PPD3, compared to 11.9% on PPD1 (*p*<0.0001) (Figure 3, A and B). This pro-inflammatory polarization remained significantly elevated on PPD7 (66.5%, *p*<0.0001) and PPD14 (58.8%, *p*<0.0001). Notably, when compared to PPD1, M2 macrophage proportions increased again at later time points, reaching 10% (p<0.01) on PPD21 and 5% on PPD28 (p<0.0001), suggesting a re-establishment of anti-inflammatory responses during later stages of uterine involution. Interestingly, we identified a distinct yet prominent macrophage subset co-expressing both M1 and M2 markers (CD86+CD206+), representing a mixed polarization phenotype. This M1/M2 subpopulation was significantly elevated on PPD1 (37.2%) compared to NP controls (7.8%, p<0.01), PPD7 (11.6%, p<0.01), and PPD14 (26.4%, p<0.05) (Figure 3, A and D). After reaching a nadir on PPD7 (11.6%), M1/M2 cells gradually increased again through PPD14 (26.4%), reaching a second peak on PPD21 (34.5%) and PPD28 (29.4%), significantly greater than both NP and PPD7 (*p*<0.05) (Figure 3, A and D). Overall, these findings point to very dynamic shifts in macrophage polarization states throughout the PP period.

### Distinct Spatial Distribution Patterns of CD206+ Macrophages in the Uterine Endometrium During Early Postpartum Uterine Remodeling

To corroborate our flow cytometry findings and to examine the localization and distribution of macrophages within the uterine parenchyma, we performed immunohistochemistry. The F4/80 marker was used to assess the total macrophage population and its distribution, while the CD206 marker was used to confirm the localization of M2 macrophages. Representative H&E histological images from all time points are shown in Figure 4A. Immunohistochemical analysis of macrophage populations was consistent with the flow cytometry findings. The total number of macrophages (F4/80+ cells) per high-power field (hpf) was markedly elevated in the early postpartum (PP) period, with PPD1 and PPD3 showing the highest numbers/hpf (52.6 and 56.1, respectively), while all subsequent PP time points exhibited lower numbers (<29) (Figure 4B). Similarly, CD206+ cells were most abundant on PPD1 (40.3), gradually decreasing on PPD3 (30), PPD7 (10.9, p < 0.01), and PPD14 (10.1, p < 0.01). A resurgence of CD206+ cells was observed on PPD21 (13.8, p < 0.01), with levels decreasing again on PPD28 (9.3, p < 0.01) (Figure 5B). Notably, on PPD1 and to a lesser extent at PPD3, we identified a large population of F4/80+ macrophages predominantly localized in the subepithelial region. These cells were no longer detected on PPD7, suggesting their transient presence during the early PP phase. Concurrently, CD206+ macrophages were largely concentrated at the myometrial-endometrial junction on PPD1, where they were primarily distributed in cell clusters (Figure 5A). Of note, CD206+ macrophages were absent from the subepithelial region at all postpartum time points. As the postpartum period progressed, they no longer formed cell clusters but remained localized closer to the myometrium (Figure 5A).

**Figure 4.**
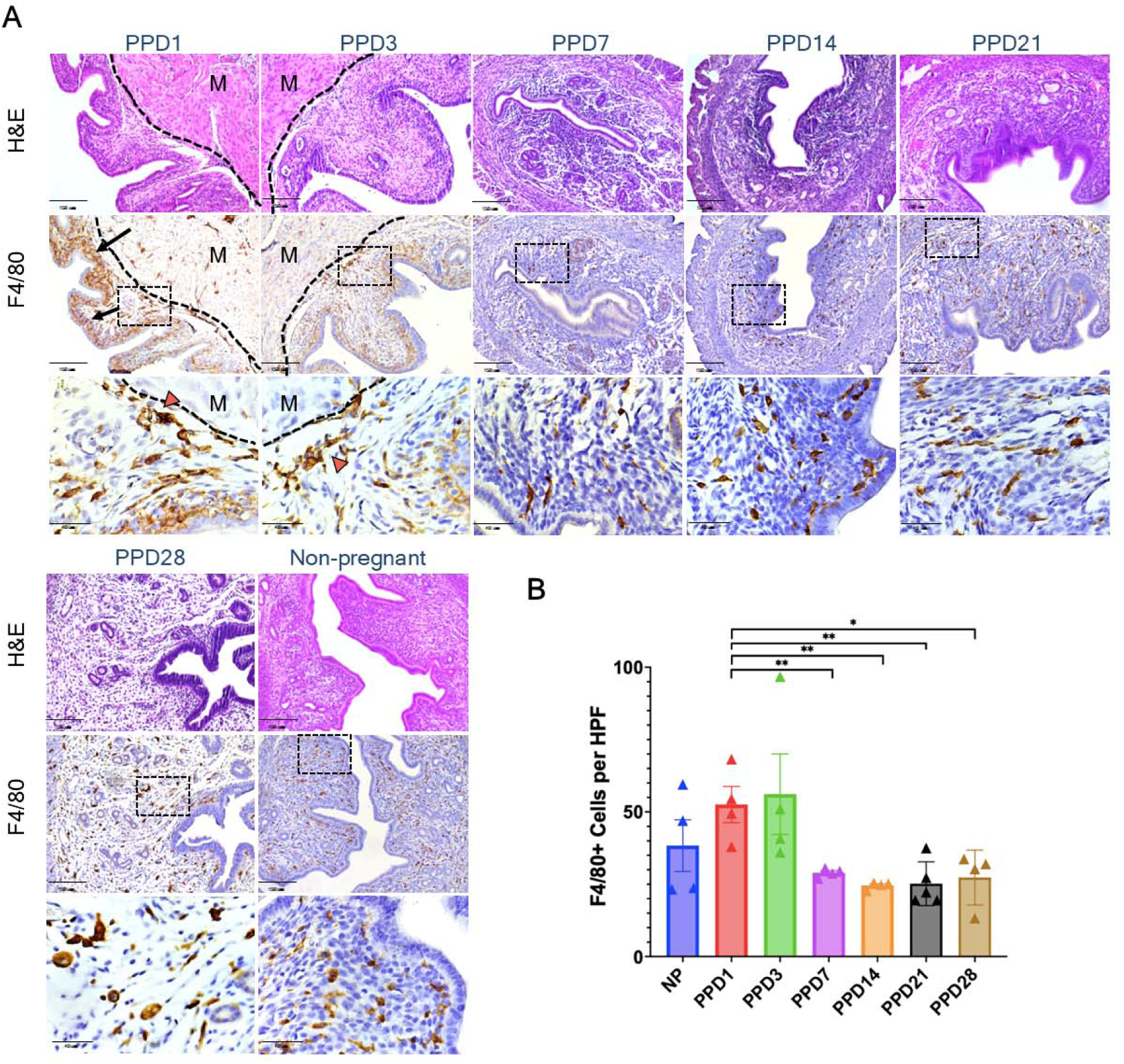
Temporal and spatial distribution of F4/80+ macrophages in the postpartum and non-pregnant uterus. (A) Representative H&E images of uterine sections at 10x magnification from non-pregnant (NP) mice and specific postpartum (PP) time points. (B) Immunohistochemical staining for F4/80+ macrophages in uterine sections from NP mice and specific PP time points, shown at low (10x, upper panel) and high (40x, lower panel) magnification. High-magnification panels correspond to regions outlined by black dashed rectangles. The dashed black line in the PPD1 and PPD3 sections delineates the endometrium–myometrium junction. Black arrows point to abundant F4/80+ macrophages localized to the subepithelial area. Red arrowheads point to F4/80+ clusters of cells localized to the endometrium–myometrium junction. (C) Quantification of F4/80+ cells per high-power field (HPF). Data is presented as mean ± SEM. M = Myometrium. *N* = 4-5 animals per group. \**p* < 0.05; **p < 0.01; \*\*\**p* < 0.001; ****p < 0.0001.

**Figure 5.**
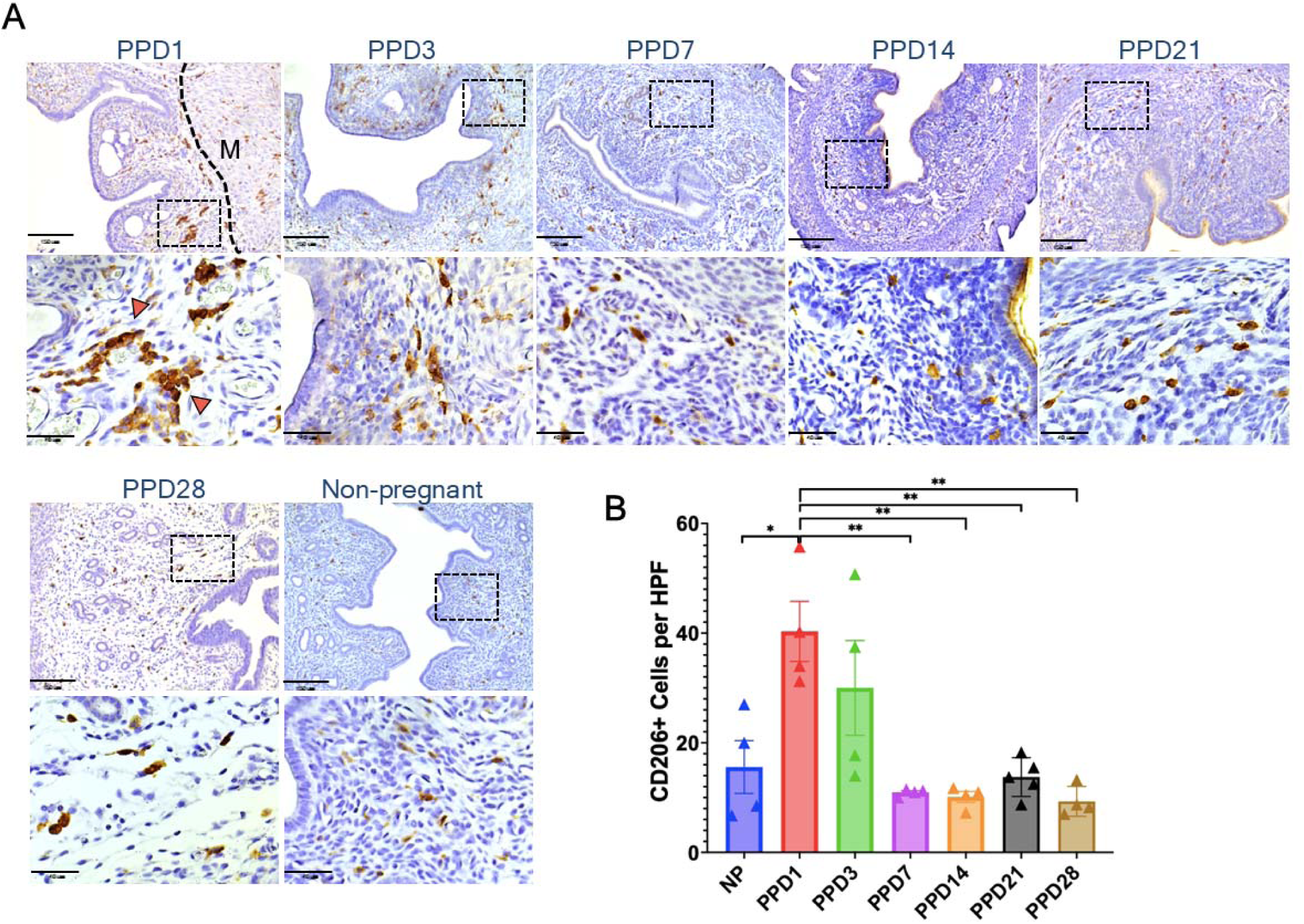
Temporal and spatial distribution of CD206+ cells in the postpartum and non-pregnant uterus. (A) Representative immunohistochemical staining for CD206+ cells in uterine tissue sections from non-pregnant (NP) and postpartum (PP) mice, shown at low (10x) and high (40x) magnification. Black arrows indicate clusters of CD206+ cells close to the endometrium– myometrium junction (noted as a black dashed line) on PPD1. (B) Quantification of CD206+ cells per high-power field (HPF). Data is presented as mean ± SEM. M = Myometrium. *N* = 4-5 animals per group. \**p* < 0.05; **p < 0.01; \*\*\**p* < 0.001; ****p < 0.0001.

### Co-localization of F4/80 and CD206 Macrophages Confirms Temporal Shifts in M2 Macrophage Populations

To further validate our flow cytometry findings and assess the spatial distribution of macrophage subpopulations in the uterine tissue, we performed immunofluorescence (IF) staining across all postpartum (PP) time points and the non-pregnant (NP) uterus. Total macrophages were identified using F4/80 immunostaining, while M2 macrophages were identified by the colocalization of CD206 with F4/80 markers. Quantification revealed trends consistent with both our flow cytometry and IHC data. Total F4/80+ macrophage numbers per high power field (hpf) were higher on PPD1, PPD3, and NP mice compared to the rest of the time points (Figure 6B). Similarly, M2 macrophages (F4/80+CD206+) absolute numbers (per hpf) were most abundant on PPD1 (26.7) and decreased on PPD3 (18.2), PPD7 (5.9, *p* < 0.01), and PPD14 (6.4, *p* < 0.01), with a secondary peak observed on PPD21 (8.3, *p* < 0.01), followed by a reduction on PPD28 (2.6, *p* < 0.001) (Figure 6C). Notably, M2 macrophages co-localizing both markers (F4/80+CD206+) were particularly evident within the cluster-forming macrophages previously observed by IHC. The proportion of CD206+ cells out of the total F4/80+ macrophages assessed by IF closely paralleled our flow cytometry findings. On postpartum PPD1, the percentage of CD206+ macrophages out of the total F4/80+ was 77.2%, representing the highest level across all time points. Compared to PPD1, this proportion declined sharply by PPD3 (46.7%, p<0.01) and continued to decrease through PPD7 and PPD14 (both 29%, p<0.0001). In NP controls, the percentage was lower (18%, p<0.0001). A secondary peak was observed on PPD21 (42.6%, p<0.001), followed by another decline on PPD28 (13.7%, p<0.0001) (Figure 6D).

**Figure 6.**
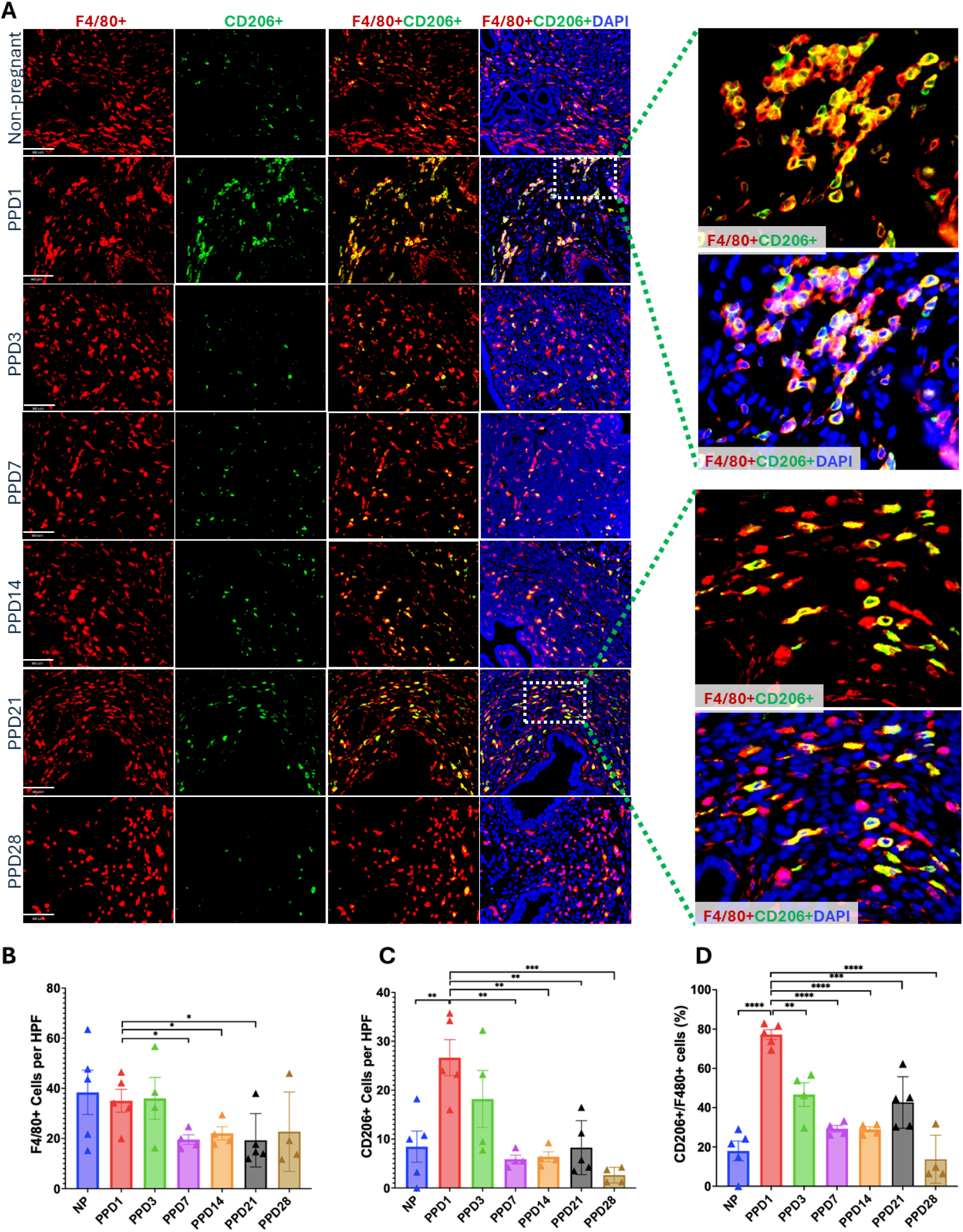
Colocalization of F4/80+ and CD206+ macrophages in the postpartum and non-pregnant uterus. (A) Representative immunofluorescence images of uterine sections from non-pregnant (NP) and postpartum (PP) mice (PPD1, PPD3, PPD7, PPD14, PPD21, PPD28), 20x magnification. Co-localization of F4/80 (red) and CD206 (green) indicates M2 macrophages; nuclei are counterstained with DAPI (blue). Higher-magnification images (40x, right panels) correspond to regions marked by dashed white boxes and show clusters of F4/80+CD206+ macrophages at PPD1 and PPD21. (B) Quantification of total F4/80+ cells per high-power field (HPF), (C) total CD206+F4/80+ cells per HPF (representing M2 macrophages), (D) and the percentage of CD206+F4/80+ cells out of total F4/80+ cells across postpartum (PP) time points and in non-pregnant (NP) control. *N* = 4-5 animals per group. \**p* < 0.05; **p < 0.01; \*\*\**p* < 0.001; ****p < 0.0001.

## DISCUSSION

Macrophages are well known to play context-specific roles in tissue remodeling, repair, and homeostasis across a variety of organs.^25–27^ These cells exhibit remarkable plasticity, adopting distinct polarization states that influence their pro-inflammatory or immunoregulatory functions depending on the local microenvironment.^28,29^ These fluctuations between different polarization states are especially relevant at the maternal– fetal interface, contributing to implantation, fetal tolerance, pregnancy maintenance, and the orchestration of parturition.^12,14,30,31^ Parturition is widely recognized as an inflammatory process characterized by elevated levels of pro-inflammatory cytokines and a substantial influx of immune cells into the decidua^32^, including macrophages.^33,34^ This recruitment has been associated with increased expression of both pro-and anti-inflammatory cytokines, including IL-1β, IL-6, IL-12, and IL-10.^35^ Consistent with this, our findings show increased leukocyte numbers in the uterus in the immediate postpartum period, including a large infiltration of macrophages.

Macrophages are the second most abundant endometrial leukocyte population and the predominant myometrial leukocyte population. During the different phases of pregnancy, macrophages undergo dynamic changes between activated M1 (proinflammatory) and alternatively activated M2 (anti-inflammatory) profiles. After coitus and in the peri-implantation period, activated M1 macrophage induce pro-inflammatory responses promoting embryo implantation. Upon trophoblast invasion and until the early phase of the second trimester, decidual macrophages display an M1/M2 profile which is thought to promote trophoblast invasion and vascular remodeling.^12^ Subsequently, after placentation is complete, the macrophages shift toward M2-dominant environment, which suppresses inflammation and prevents fetal rejection. During parturition, which is considered a pro-inflammatory event, M1 macrophages predominate again.^33^ However, there is little evidence regarding macrophage polarization profiles during the postpartum period. Egari et al. showed infiltration of arginase-expressing M2-like macrophages in the postpartum uterus which peaked on PPD3 and subsequently diminished, but their investigation did not extend beyond PPD7.^36^

In our study, we observed an early dominance of CD206+ (M2-like) macrophages on PPD1, gradually shifting toward a CD86+ dominant (M1-like) profile by PPD7–PPD14, suggesting an orchestrated transition aimed at balancing tissue repair and inflammation. The early transition of the endometrium toward an M2-dominant anti-inflammatory environment is likely necessary to achieve wound healing, including epithelial repair, and facilitating rapid tissue barrier restoration and wound closure, functions previously associated with M2-like cells in wound healing of different tissues.^17,37,38^ Indeed, abnormally increased M1 polarization associated with elevated IFNγ in the postpartum was noted in RBPJ-deficient mice, which exhibited defective postpartum uterine repair.^39^ As macrophages shift toward an M1-like profile, they likely contribute to host defense from pathogens and support healthy tissue remodeling. This response may also help prevent excessive M2-like activation, which has been associated with fibrotic outcomes.^40^

Interestingly, our findings reveal a previously unreported secondary peak in M2-like macrophage subset on PPD21, the significance of which remains to be fully elucidated. This late increase may reflect a second wave of tissue remodeling, potentially associated with the final restoration of endometrial function. For example, macrophages are crucial for the postpartum clearance of senescent cells from the implantation scar, a gradual process of tissue repair that takes approximately a month to complete in mice.^14^ Elucidating the functional role of this polarization shift is important, as it may mark a vulnerable postpartum window in which disruptions in macrophage-mediated repair could impair endometrial recovery or predispose to pathological conditions. One possible explanation for this late rise in M2-like macrophages is its association with physiological changes linked to lactation and weaning. In the mammary gland, macrophages are essential for normal involution following weaning, with their numbers rising markedly during the irreversible phase of involution. At this stage, they adopt an M2-like, tissue-repair phenotype that facilitates epithelial cell death, alveolar regression, and adipocyte repopulation.^41–44^ A similar process may occur in the uterus, whereby M2-like macrophages increase again in response to systemic or local cues associated with the cessation of lactation. In our study, uteri analyzed at earlier postpartum time points (PPD7 and PPD14) may not have exhibited this M2 increase because their pups were still actively nursing. In contrast, by PPD21, coinciding with the typical weaning period when pups begin consuming solid food, mice may undergo hormonal and immunological shifts that promote M2 polarization. These observations raise the intriguing possibility that the timing of pup weaning and growth could influence uterine immune dynamics during late postpartum remodeling.

One of the most notable findings of this study was the presence of a macrophage subset co-expressing both M1-and M2-like surface markers. These CD86+CD206+ cells were particularly elevated on PPD1-PPD3, with a secondary peak observed on PPD21 and PPD28. To our knowledge, these cells have not been previously characterized in the postpartum uterus. The literature offers differing interpretations regarding cells expressing both M1 and M2 markers. Some studies suggest that co-expression reflects the broad spectrum of macrophage activation and their inherent functional plasticity, representing transitional states between classical and alternative polarization.^45,46^ Others propose that these cells are mature macrophages with a stable, mixed phenotype, adapted to perform complex roles in specific tissues or physiological contexts.^38,47^ Another theory suggests that macrophages may undergo sequential reprogramming, retaining markers from prior activation states as they transition in response to new environmental cues. This co-expression could reflect the dynamic reprogramming of these cells.^20,45^ Additionally, some studies have linked the presence of macrophages with mixed M1/M2 phenotypes to disease-associated dysregulation. In pathological contexts such as cancer, fibrosis, or chronic inflammation, these hybrid macrophages may arise from prolonged exposure to conflicting signals. Such hybrid macrophages may reflect a maladaptive response that contributes to disease progression by sustaining inflammation or excessively promoting tissue remodeling.^47–49^

Further studies are needed to determine the functional role of these M1/M2 cells in the postpartum remodeling process. In this context, surface marker-based phenotyping may oversimplify the diverse activation states of macrophages.^18,38,45^ To overcome this limitation, future work will incorporate single-cell RNA sequencing (scRNA-seq) to provide higher-resolution insights into macrophage polarization and functional heterogeneity beyond the conventional M1/M2 framework to better define their roles in the postpartum uterine remodeling.

Our immunohistochemical results revealed distinct spatial and temporal patterns of macrophage distribution in the postpartum uterus, underscoring their heterogeneity and dynamic roles in tissue remodeling. The transient accumulation of F4/80+ macrophages in the subepithelial region on PPD1, and to a lesser extent on PPD3, suggests an early infiltration wave likely triggered by parturition-associated epithelial disruption, providing both antimicrobial defense and support for initial tissue repair. Their absence by PPD7 may reflect resolution of this early response through macrophage-mediated clearance of debris and restoration of epithelial integrity.^14^ In contrast, CD206+ cells were consistently localized at the myometrial–endometrial junction, often forming clusters of cells in early postpartum, suggesting a more sustained and spatially restricted role for M2-polarized macrophages. Such positioning likely reflects local cues that direct their involvement in processes such as neovascularization and smooth muscle regeneration.^28,37,38^ Taken together, these findings are consistent with prior reports showing that macrophages first localize beneath the endometrial lining and later spread into the endometrium and myometrium, where they contribute to collagen resorption and tissue remodeling.^13,30,50^ These spatial dynamics highlight the influence of uterine microenvironments on macrophage function and suggest that distinct subsets act in a temporally and regionally coordinated manner to promote postpartum healing.

In conclusion, we characterized the dynamics of macrophage polarization in the uterus throughout the postpartum period, demonstrating rapid changes in macrophage polarization profiles and subsets in the process of uterine tissue remodeling and repair. Additional studies are needed to better define the functions and mechanisms of various macrophage subsets in postpartum uterine remodeling. Such deeper understanding may pave the way for macrophage-targeted therapies that promote healthy scarless endometrial regeneration.

## ACKNOWLEDGEMENTS

This work was supported by Eunice Kennedy Shriver National Institute of Child Health and Human Development (NICHD) grants 1RO1HD10926-01A1 (to RT) and 3RO1HD10926-01S1 (to RT). We appreciate the use of the Yale Flow Cytometry Core Facility. We are grateful to Lesley Devine (Yale Flow Cytometry Core) for her advice and technical help. We thank Amos Brooks (Department of Pathology, Yale School of Medicine) for his helpful advice.

## COMPETING INTERESTS

The authors declare that they have no competing interests.

## Notes

### Competing Interest Statement

The authors have declared no competing interest.

## REFERENCES

1. Suarez AC, Gimenez CJ, Russell SR, et al. Pregnancy-induced remodeling of the murine reproductive tract: a longitudinal in vivo magnetic resonance imaging study. Sci Rep. 2024;14(1):586. doi:10.1038/s41598-023-50437-1

2. Spooner MK, Lenis YY, Watson R, Jaimes D, Patterson AL. The role of stem cells in uterine involution. Reproduction. 2021;161(3):R61–R77. doi:10.1530/REP-20-0425

3. Dai T, Ma Z, Guo X, et al. Study on the Pattern of Postpartum Uterine Involution in Dairy Cows. Animals. 2023;13(23):3693. doi:10.3390/ani13233693

4. Baah-Dwomoh A, McGuire J, Tan T, De Vita R. Mechanical Properties of Female Reproductive Organs and Supporting Connective Tissues: A Review of the Current State of Knowledge. Appl Mech Rev. 2016;68(060801). doi:10.1115/1.4034442

5. Nguyen TTTN, Shynlova O, Lye SJ. Matrix Metalloproteinase Expression in the Rat Myometrium During Pregnancy, Term Labor, and Postpartum1. Biol Reprod. 2016;95(1):24, 1–14. doi:10.1095/biolreprod.115.138248

6. Ryvnyak VV, Dulgieru OF. Elastase Involvement in Extracellular and Intracellular Collagen Degradation during Postpartum Involution of the Uterus. Bull Exp Biol Med. 2003;136(2):206–208. doi:10.1023/A:1026395613444

7. Ryvnyak VV, Gudumak VS, Rybakova MA, Grumeza OF, Pelin AV. Extra-and intracellular collagen resorption by smooth muscle cells in postpartum uterine involution. Bull Exp Biol Med. 1999;127(1):96–98. doi:10.1007/BF02432812

8. Dessouky DA. Myometrial changes in postpartum uterine involution. Am J Obstet Gynecol. 1971;110(3):318–329. doi:10.1016/0002-9378(71)90721-6

9. Dadarwal D, Palmer C, Griebel P. Mucosal immunity of the postpartum bovine genital tract. Theriogenology. 2017;104:62–71. doi:10.1016/j.theriogenology.2017.08.010

10. Machado VS, Silva TH. Adaptive immunity in the postpartum uterus: Potential use of vaccines to control metritis. Theriogenology. 2020;150:201–209. doi:10.1016/j.theriogenology.2020.01.040

11. Azawi OI. Postpartum uterine infection in cattle. Anim Reprod Sci. 2008;105(3):187–208. doi:10.1016/j.anireprosci.2008.01.010

12. Zhang YH, He M, Wang Y, Liao AH. Modulators of the Balance between M1 and M2 Macrophages during Pregnancy. Front Immunol. 2017;8:120. doi:10.3389/fimmu.2017.00120

13. Parakkal PF. Macrophages: The time course and sequence of their distribution in the postpartum uterus. J Ultrastruct Res. 1972;40(3):284–291. doi:10.1016/S0022-5320(72)90101-3

14. Egashira M, Hirota Y, Shimizu-Hirota R, et al. F4/80+ Macrophages Contribute to Clearance of Senescent Cells in the Mouse Postpartum Uterus. Endocrinology. 2017;158(7):2344–2353. doi:10.1210/en.2016-1886

15. Mills CD, Kincaid K, Alt JM, Heilman MJ, Hill AM. M-1/M-2 Macrophages and the Th1/Th2 Paradigm1. J Immunol. 2000;164(12):6166–6173. doi:10.4049/jimmunol.164.12.6166

16. Mantovani A, Sica A, Sozzani S, Allavena P, Vecchi A, Locati M. The chemokine system in diverse forms of macrophage activation and polarization. Trends Immunol. 2004;25(12):677–686. doi:10.1016/j.it.2004.09.015

17. Novak ML, Koh TJ. Macrophage phenotypes during tissue repair. J Leukoc Biol. 2013;93(6):875–881. doi:10.1189/jlb.1012512

18. Strizova Z, Benesova I, Bartolini R, et al. M1/M2 macrophages and their overlaps – myth or reality? Clin Sci. 2023;137(15):1067–1093. doi:10.1042/CS20220531

19. Orecchioni M, Ghosheh Y, Pramod AB, Ley K. Macrophage Polarization: Different Gene Signatures in M1(LPS+) vs. Classically and M2(LPS–) vs. Alternatively Activated Macrophages. Front Immunol. 2019;10. doi:10.3389/fimmu.2019.01084

20. Liu SX, Gustafson HH, Jackson DL, Pun SH, Trapnell C. Trajectory analysis quantifies transcriptional plasticity during macrophage polarization. Sci Rep. 2020;10(1):12273. doi:10.1038/s41598-020-68766-w

21. Tal R, Liu Y, Pluchino N, Shaikh S, Mamillapalli R, Taylor HS. A Murine 5-Fluorouracil-Based Submyeloablation Model for the Study of Bone Marrow-Derived Cell Trafficking in Reproduction. Endocrinology. 2016;157(10):3749–3759. doi:10.1210/en.2016-1418

22. Zhu X, Chen S, Zhang P, et al. Granulocyte-macrophage colony-stimulating factor promotes endometrial repair after injury by regulating macrophages in mice. J Reprod Immunol. 2023;160:104156. doi:10.1016/j.jri.2023.104156

23. Duan J, Liu X, Wang H, Guo SW. The M2a macrophage subset may be critically involved in the fibrogenesis of endometriosis in mice. Reprod Biomed Online. 2018;37(3):254–268. doi:10.1016/j.rbmo.2018.05.017

24. Lv H, Sun H, Wang L, et al. Targeting CD301 + macrophages inhibits endometrial fibrosis and improves pregnancy outcome. EMBO Mol Med. 2023;15(9):e17601. doi:10.15252/emmm.202317601

25. Orvalho JM, Fernandes JCH, Moraes Castilho R, Fernandes GVO. The Macrophage’s Role on Bone Remodeling and Osteogenesis: a Systematic Review. Clin Rev Bone Miner Metab. 2023;21(1):1–13. doi:10.1007/s12018-023-09286-9

26. Michalski MN, McCauley LK. Macrophages and skeletal health. Pharmacol Ther. 2017;174:43–54. doi:10.1016/j.pharmthera.2017.02.017

27. O’Rourke SA, Dunne A, Monaghan MG. The Role of Macrophages in the Infarcted Myocardium: Orchestrators of ECM Remodeling. Front Cardiovasc Med. 2019;6. doi:10.3389/fcvm.2019.00101

28. Mantovani A, Biswas SK, Galdiero MR, Sica A, Locati M. Macrophage plasticity and polarization in tissue repair and remodelling. J Pathol. 2013;229(2):176–185. doi:10.1002/path.4133

29. Wynn TA, Vannella KM. Macrophages in Tissue Repair, Regeneration, and Fibrosis. Immunity. 2016;44(3):450–462. doi:10.1016/j.immuni.2016.02.015

30. Mackler AM, Iezza G, Akin MR, McMillan P, Yellon SM. Macrophage Trafficking in the Uterus and Cervix Precedes Parturition in the Mouse. Biol Reprod. 1999;61(4):879–883. doi:10.1095/biolreprod61.4.879

31. Lv M, Jia Y, Dong J, Wu S, Ying H. The landscape of decidual immune cells at the maternal–fetal interface in parturition and preterm birth. Inflamm Res. 2025;74(1):44. doi:10.1007/s00011-025-02015-6

32. Osman I, Young A, Ledingham MA, et al. Leukocyte density and pro inflammatory cytokine expression in human fetal membranes, decidua, cervix and myometrium before and during labour at term. Mol Hum Reprod. 2003;9(1):41–45. doi:10.1093/molehr/gag001

33. Hamilton S, Oomomian Y, Stephen G, et al. Macrophages Infiltrate the Human and Rat Decidua During Term and Preterm Labor: Evidence That Decidual Inflammation Precedes Labor1. Biol Reprod. 2012;86(2):39, 1-9. doi:10.1095/biolreprod.111.095505

34. Tal R, Kisa J, Abuwala N, et al. Bone Marrow-Derived Progenitor Cells Contribute to Remodeling of the Postpartum Uterus. Stem Cells. 2021;39(11):1489–1505. doi:10.1002/stem.3431

35. Brown MB, von Chamier M, Allam AB, Reyes L. M1/M2 Macrophage Polarity in Normal and Complicated Pregnancy. Front Immunol. 2014;5. doi:10.3389/fimmu.2014.00606

36. Yoshii A, Kitahara S, Ueta H, Matsuno K, Ezaki T. Role of Uterine Contraction in Regeneration of the Murine Postpartum Endometrium1. Biol Reprod. 2014;91(2):32, 1-10. doi:10.1095/biolreprod.114.117929

37. Hesketh M, Sahin KB, West ZE, Murray RZ. Macrophage Phenotypes Regulate Scar Formation and Chronic Wound Healing. Int J Mol Sci. 2017;18(7):1545. doi:10.3390/ijms18071545

38. Yan L, Wang J, Cai X, et al. Macrophage plasticity: signaling pathways, tissue repair, and regeneration. MedComm. 2024;5(8):e658. doi:10.1002/mco2.658

39. Strug MR, Su RW, Kim TH, et al. RBPJ mediates uterine repair in the mouse and is reduced in women with recurrent pregnancy loss. FASEB J. 2018;32(5):2452–2466. doi:10.1096/fj.201701032R

40. Murray PJ, Wynn TA. Protective and pathogenic functions of macrophage subsets. Nat Rev Immunol. 2011;11(11):723–737. doi:10.1038/nri3073

41. Martinson HA, Jindal S, Durand-Rougely C, Borges VF, Schedin P. Wound healing-like immune program facilitates postpartum mammary gland involution and tumor progression. Int J Cancer. 2015;136(8):1803–1813. doi:10.1002/ijc.29181

42. O’Brien J, Martinson H, Durand-Rougely C, Schedin P. Macrophages are crucial for epithelial cell death and adipocyte repopulation during mammary gland involution. Development. 2012;139(2):269–275. doi:10.1242/dev.071696

43. Li Y, Pang Z, Dong X, et al. MUC1 induces M2 type macrophage influx during postpartum mammary gland involution and triggers breast cancer. Oncotarget. 2017;9(3):3446–3458. doi:10.18632/oncotarget.23316

44. Stewart TA, Hughes K, Hume DA, Davis FM. Developmental Stage-Specific Distribution of Macrophages in Mouse Mammary Gland. Front Cell Dev Biol. 2019;7. doi:10.3389/fcell.2019.00250

45. Smith TD, Tse MJ, Read EL, Liu WF. Regulation of macrophage polarization and plasticity by complex activation signals. Integr Biol Quant Biosci Nano Macro. 2016;8(9):946–955. doi:10.1039/c6ib00105j

46. Herb M, Schatz V, Hadrian K, et al. Macrophage variants in laboratory research: most are well done, but some are RAW. Front Cell Infect Microbiol. 2024;14. doi:10.3389/fcimb.2024.1457323

47. Italiani P, Boraschi D. From Monocytes to M1/M2 Macrophages: Phenotypical vs. Functional Differentiation. Front Immunol. 2014;5. doi:10.3389/fimmu.2014.00514

48. Xue JD, Gao J, Tang AF, Feng C. Shaping the immune landscape: Multidimensional environmental stimuli refine macrophage polarization and foster revolutionary approaches in tissue regeneration. Heliyon. 2024;10(17):e37192. doi:10.1016/j.heliyon.2024.e37192

49. Furgiuele S, Descamps G, Cascarano L, et al. Dealing with Macrophage Plasticity to Address Therapeutic Challenges in Head and Neck Cancers. Int J Mol Sci. 2022;23(12):6385. doi:10.3390/ijms23126385

50. Mackler AM, Green LM, McMillan PJ, Yellon SM. Distribution and Activation of Uterine Mononuclear Phagocytes in Peripartum Endometrium and Myometrium of the Mouse. Biol Reprod. 2000;62(5):1193–1200. doi:10.1095/biolreprod62.5.1193

