## Supplemental Table 1 for "Dynamics of Macrophage Polarization Reveal Unique Phenotypes During Mouse Postpartum Uterine Remodeling"

**Supplementary Table S1.** Antibodies used in flow cytometry, immunohistochemistry, and immunofluorescence.

| Antibody | Manufacturer | Identifier | Concentration |
| --- | --- | --- | --- |
| Goat polyclonal anti-CD206 | R&D Systems | AF2535 | 1:400 |
| Rabbit monoclonal anti-F4/80 | Abcam | ab300421 | 1:500 |
| Donkey anti-rabbit Alexa Fluor 594 | ThermoFisher Scientific | A21207 | 1:200 |
| Donkey anti-goat Alexa Fluor 488 | ThermoFisher Scientific | A11055 | 1:200 |
| Goat anti-rabbit biotinylated antibody | Vector Laboratories | 30014 | 1:200 |
| Rabbit anti-goat biotinylated antibody | Vector Laboratories | 30056 | 1:200 |
| FITC anti-mouse CD3 | Biolegend | B388790 | 1:100 |
| APC/Cy7 anti-mouse CD11b | Biolegend | B421065 | 1:100 |
| BV605 anti-mouse CD45 | Biolegend | B396771 | 1:100 |
| Pacific Blue anti-mouse CD86 | Biolegend | B377787 | 1:100 |
| AF700 anti-mouse CD11C | Biolegend | B418344 | 1:100 |
| BV711 anti-mouse F4/80 | Biolegend | B423715 | 1:100 |
| PE anti-mouse Ly6G | Biolegend | B397726 | 1:100 |
| APC anti-mouse CD206 | Biolegend | B411653 | 1:100 |
| PerCP/Cyanine5.5 anti-mouse 301b | Biolegend | B392244 | 1:100 |
| PE-Cy7 anti-mouse CD163 | Biolegend | B407884 | 1:100 |
